# A meta-analysis of clinical cases of reversion mutations in *BRCA* genes identifies signatures of DNA end-joining repair mechanisms driving therapy resistance

**DOI:** 10.1101/2020.07.22.215673

**Authors:** Luis Tobalina, Joshua Armenia, Elsa Irving, Mark J. O’Connor, Josep V. Forment

## Abstract

Germline mutations in the *BRCA1* or *BRCA2* genes predispose to hereditary breast and ovarian cancer and, mostly in the case of *BRCA2*, are also prevalent in cases of pancreatic and prostate malignancies. Tumours from these patients tend to lose both copies of the wild type *BRCA* gene, which makes them exquisitely sensitive to platinum drugs and PARP inhibitors (PARPi), treatments of choice in these disease settings. Reversion secondary mutations with the capacity of restoring BRCA protein expression have been documented in the literature as *bona fide* mechanisms of resistance to these treatments. Here, we perform a detailed analysis of clinical cases of reversion mutations described in *BRCA1* and *BRCA2*, which underlines the different importance of BRCA protein domains in contributing to resistance and the potential key role of mutagenic end-joining DNA repair pathways in generating reversions. Our analyses suggest that pharmacological inhibition of these repair pathways could improve durability of drug treatments and highlights potential interventions to both prevent the appearance of reversions and provide new therapeutic opportunities after their acquisition.

**Highlights:**

- Comprehensive analysis of reversion mutations in *BRCA* genes identified in clinical cases of resistance to platinum or PARPi
- Revertant proteins devoid of parts of the original sequence, identifying key protein functions involved in resistance
- Hypomorph revertant BRCA proteins suggest potential new therapeutic opportunities to overcome resistance
- Prevalence of mutational end-joining DNA repair mechanisms leading to reversions, especially in those affecting *BRCA2*
- Pharmacological inhibition of mutational end-joining DNA repair could improve durability of drug treatments

## Introduction

Mutations in the breast cancer susceptibility genes *BRCA1* and *BRCA2* have been associated since the 1990s to hereditary cases of breast and ovarian cancers. Patients with inactivating germline mutations in these genes – usually, small insertion-deletions (INDELs) or single-nucleotide variants (SNV) causing frameshifts in the open reading frame (ORF) and premature STOP codons - have an increased risk of developing breast cancers and, to a lesser extent, ovarian cancers. Following the classical ‘two-hit’ model of tumour suppressor inactivation, tumours from these patients tend to lose functionality of the remaining *BRCA* wild type allele, usually by loss of heterozygosity[1]. Loss of *BRCA* genes fosters genomic instability in tumours due to the key role BRCA1 and BRCA2 proteins play in DNA replication-fork protection and homologous recombination repair (HRR), a high-fidelity DNA repair pathway involved in the repair of DNA double-strand breaks and other genotoxic lesions[2].

Although loss of BRCA function could be beneficial for tumour development, it also makes tumour cells exquisitely sensitive to DNA crosslinkers such as platinum drugs. DNA crosslinks, particularly the ones formed in between the two DNA strands, rely on HRR for their efficient repair, explaining why platinum-based therapies have proven beneficial in *BRCA* mutant patients[3]. More recently, it was discovered that inhibition of poly(ADP-ribose) polymerase (PARP) shows synthetic lethality with BRCA deficiency, again linked to HRR defects[4, 5]. PARP inhibitors (PARPi) have been developed clinically and are now approved in breast, pancreatic, prostate and ovarian settings[6–8].

Platinum or PARPi treatments are effective in tumours with BRCA mutations but resistance arises. Although several different mechanisms of platinum and PARPi resistance have been described pre-clinically[2, 9], acquisition of secondary mutations in *BRCA* genes restoring the open reading frame and detected upon treatment progression is the only resistance mechanism currently validated in the clinic, with the first examples described more than a decade ago[10–12]. In this study, we analyse clinical examples of secondary mutations acquired in *BRCA* genes described in the literature[10–39] to gain insights into the mutational mechanisms driving their acquisition, and the importance of the different BRCA protein domains in generating resistance to drug treatment.

## Results

### Patients progressing on platinum or PARPi treatment accumulate reversion mutations in BRCA genes

We analysed sequencing data available in the literature from tumour or ctDNA paired samples collected from 327 patients with mutations in *BRCA1* or *BRCA2* on progression after platinum or PARPi treatment (**Supp Table S1**). This patient cohort is heavily biased towards ovarian cancer (234/327 patients), explained by germline mutations in *BRCA* genes being prevalent in this tumour type[40], and by platinum and PARPi being approved standard of care therapies in that disease setting[41] (**Fig 1a**). Response to PARPi in this cohort is predominantly for olaparib (93/241 patients) and rucaparib (73/241 patients; **Supp Table S2**).

**Figure 1.**
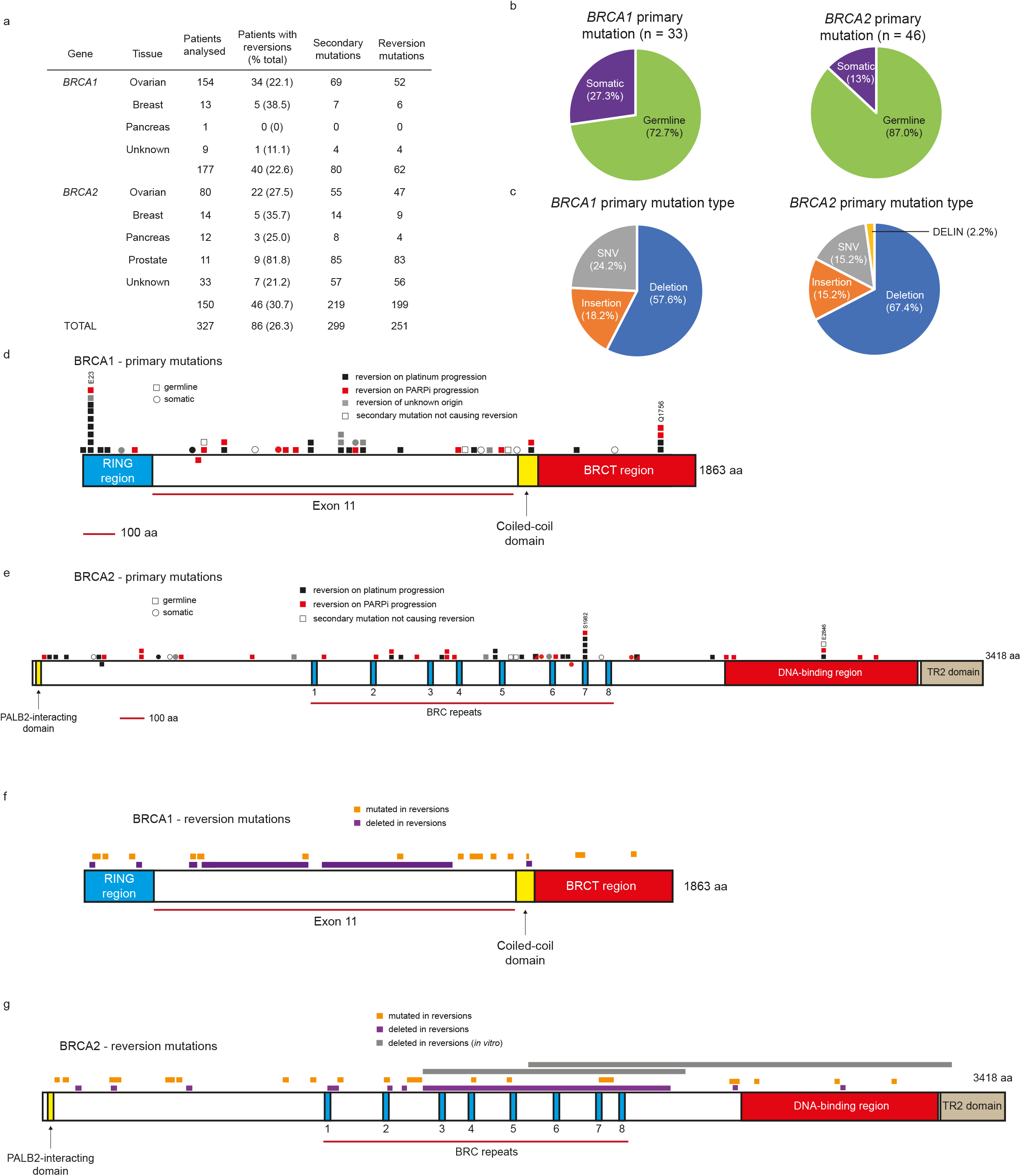
**(a)** Patients analysed in this study and number of secondary mutations and reversions identified. **(b)** Origin of primary mutations. **(c)** Type of primary mutations. SNV: single-nucleotide variant. DELIN: deletion + insertion. **(d)** Distribution of secondary mutations on the domain structure of BRCA1 and their outcome (reversion vs no reversion) depending on type of treatment received and position of the primary mutation. Each square/dot represents a patient. **(e)** Same as in **d** but for BRCA2. **(f)** Position and outcome (mutation vs deletion) of reversion mutations identified in this patient cohort on the domain structure of BRCA1. **(g)** Same as in **f** but for BRCA2. Reversions identified *in vitro* are depicted in grey for comparative purposes.

299 secondary mutations on *BRCA* genes on progression after treatment with platinum or PARPi were identified in 99 patients (46 carrying primary *BRCA1* mutations; 53 carrying primary *BRCA2* mutations; **Supp Table S1**). 269 of these secondary mutations (70 detected in 40 *BRCA1* mutant patients; 199 in 46 *BRCA2* mutant patients) corrected the original mutation in the tumour or re-established the *BRCA* open reading frame that was disrupted by it (**Supp Table S1**). As a consequence, these secondary mutations had the potential to restore BRCA protein expression, and we will refer to them as reversions (**Supp Table S2**). The percentage of *BRCA* patients accumulating reversion mutations on progression after platinum or PARPi across the different tumour types analysed was 26.3%. The percentage of patients with detectable reversions in *BRCA1* was lower than of those in *BRCA2* (22.6% vs 30.7%), although this was not statistically significant (two-tailed two-proportions test *p* value 0.12) (**Fig 1a**).

Primary mutations in *BRCA* genes in this patient cohort are predominantly from germline origin (24/33 cases for *BRCA1*; 40/46 for *BRCA2*; **Fig 1b; Supp Table S3**). They are mostly insertions or deletions (25/33 cases for *BRCA1*; 39/46 for *BRCA2*) causing frameshifts leading to premature STOP codon gains (23/25 cases for *BRCA1*; 38/39 for *BRCA2*; **Fig. 1c; Supp Table S3**). They are preferentially located in the hot spot mutated regions encoding the RING or BRCT domains of the BRCA1 protein or in the sequence comprising exon 11 of the gene, and for BRCA2, in the regions encoding the BRC repeats or the N-terminal part of the protein, between the PALB2-interacting domain and the BRC repeats[40]. Particularly well represented are the Ashkenazi Jew founder germline mutations[42] *BRCA1* 185delAG (c.68_69delAG; E23Vfs*; 9 cases) and 5382insC (c.5266insC; Q1756Pfs*; 4 cases) (**Fig 1d**), and *BRCA2* 6174delT (c.5946delT; S1982Rfs*; 5 cases) (**Fig. 1e**). As expected, all reversions occurring in tumours where the primary mutation caused a frameshift consisted of a secondary insertion or deletion restoring the ORF. Reversions where the primary mutation was a SNV occurred through a secondary mutation involving another SNV (17/23 *BRCA1* cases; 6/14 in *BRCA2*), but also through deletions (6/23 *BRCA1* cases; 8/14 in *BRCA2*) (**Supp Table S3**). For SNVs, the nature of the reversion mutation seemed to be dependent on the genomic location of the primary mutation (see below). The type of treatment received did not seem to result in secondary mutations and reversions being preferentially accumulated in specific regions of either BRCA1 or BRCA2 proteins, although the limited number of patients analysed brings caution to the interpretation of the data (**Fig 1d,e**).

### Importance of the different BRCA protein domains in conferring resistance to platinum or PARPi

Mapping of the consequences of reversion mutations on the BRCA proteins shed light on the functional importance of the different protein domains to confer resistance to treatment. In BRCA1, reversion mutations in the exon 11 region resulted in deletions of considerable amino acid length (**Fig 1f**). Two extreme cases resulted in deletions of 860 bp (286 aa) and 1170 bp (389 aa) in patients with ovarian cancer progressing on platinum or PARPi, respectively[24, 25] (**Supp Table S2**). This suggests that most of the protein region encoded by exon 11 is dispensable with regards to the BRCA1 function required to confer resistance to platinum or PARPi, most likely HRR. These data are in agreement with studies performed *in vitro*, where expression of a BRCA1 protein hypomorph lacking most or the entire sequence encoded by *BRCA1* exon 11 was shown to confer resistance to these drugs[43]. On the contrary, reversions on the RING, coiled-coil and BRCT domains of BRCA1 resulted in smaller deletions or mutations in the amino acid sequence (**Fig 1f**), suggesting that there is less flexibility to the scale of amino acid changes that can take place in these regions without fundamentally affecting BRCA1 function. Importantly, the RING domain is required for BRCA1 protein stability and ubiquitin E3 ligase activity through its interaction with BARD1[44], while the BRCT domains are also required for stability and protein-protein interactions[45, 46]. It will be important to determine experimentally the functional consequences of such deletions and mutations on BRCA1 function in HRR to fully understand what the minimum requirements for a revertant protein are to produce resistance to platinum drugs or PARPi.

Similar to the *BRCA1* cases, significant deletions affected the region encoded by exon 11 in *BRCA2* reversions. This region encodes the BRC repeats, which are the binding domains for the RAD51 recombinase, essential for the BRCA2 function in HRR[47](**Fig 1g**). One extreme case of such deletions (2541 bp) in a breast cancer patient not responding to olaparib resulted in the deletion of 6 of the 8 BRC repeats[16] (**Supp Table S2**). No reversion, however, has been described resulting in the complete elimination of all BRC repeats, suggesting that there is a minimum of at least two of these required to preserve BRCA2 function in conferring resistance to platinum or PARPi[48]. In fact, mice carrying homozygous deletion of *BRCA2* exon 11 are inviable[49]. The N-terminal or the DNA-binding region of BRCA2 only accumulated reversions with smaller deletions or mutations, suggesting that there are more constraints around amino acid changes in these regions, despite the fact that *in vitro* some reversion events resulted in the complete removal of the DNA-binding region of BRCA2[10] (**Fig 1g; Supp Table S4**).

### Different types of secondary mutations in BRCA1 and BRCA2 patients

Secondary mutations were classified as pure insertions or deletions (either of 1 bp or more), SNVs or cases where both insertions and deletions occurred (named for convenience as DELINs). Distribution of the type of secondary mutations observed in *BRCA* genes differed between *BRCA1* and *BRCA2* in this cohort. While deletions accounted for most of the secondary mutations detected in both genes, prevalence of these in *BRCA2* mutant patients was higher (77.6% vs 58.8% in *BRCA1*, two-tailed two-proportions test *p* value 0.001), especially of deletions of more than 1 bp (37.5% in *BRCA1* vs 68.0% in *BRCA2,* two-tailed two-proportions test *p* value 3.56E-06; **Fig 2a**). This is despite the fact that prevalence of primary mutations in exon 11, the most flexible region to accommodate amino acid changes in *BRCA* genes (see above), is similar between both genes in this patient cohort (55.1% in *BRCA1* vs 54.7% in *BRCA2,* two-tailed two-proportions test *p* value NS; **Supp Table S3**). The type of treatment after which the secondary mutations were detected did not seem to affect the type of mutation acquired except for deletions of 1 bp that were enriched in *BRCA2* after PARPi treatment (1.5% after platinum vs 15.2% after PARPi, two-tailed two-proportions test *p* value 0.007; **Fig 2b,c; Supp Table S2**). The limited number of cases, however, warrants caution when analysing these results.

**Figure 2.**
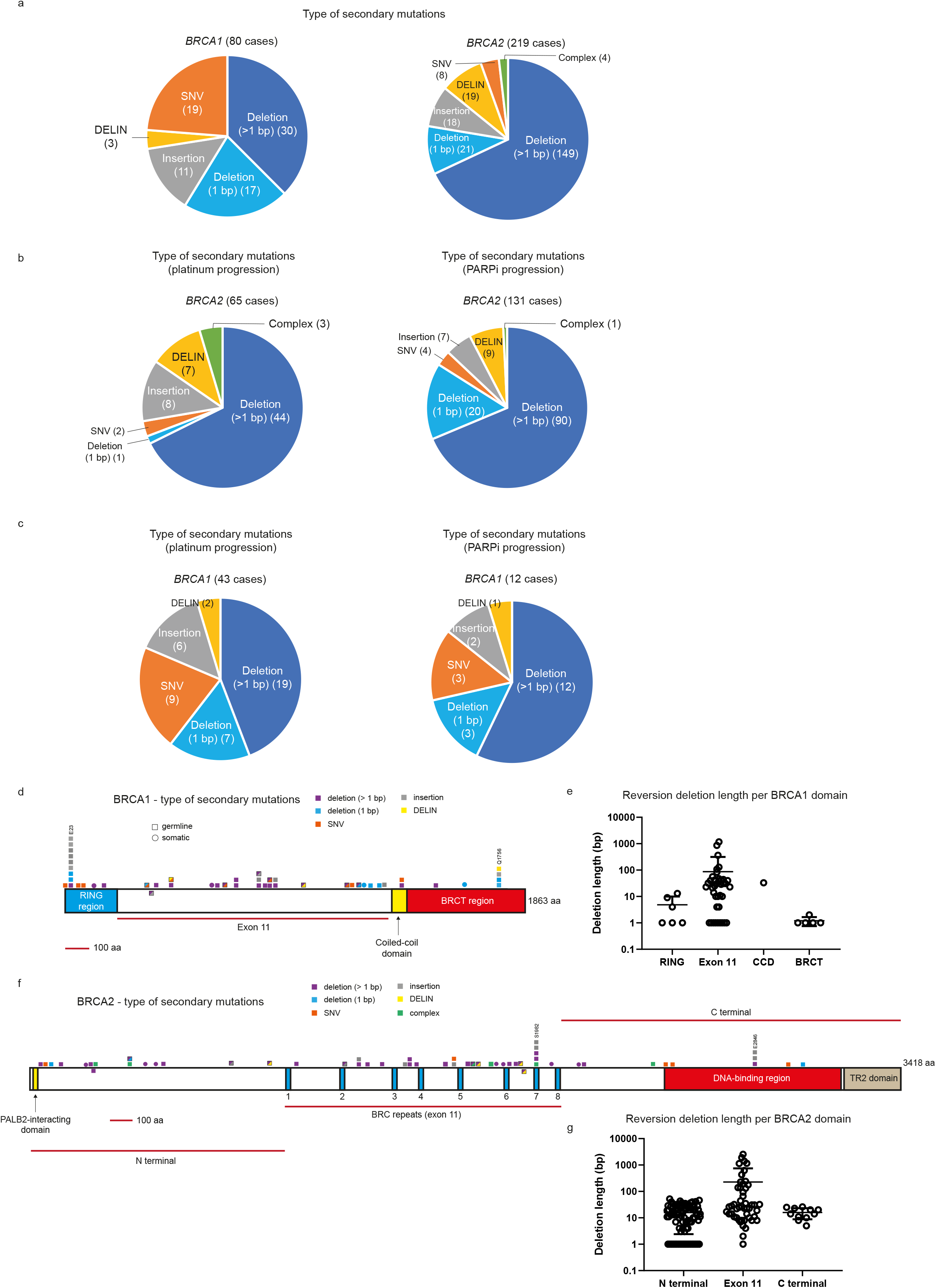
**(a)** Distribution of type of secondary mutations identified in *BRCA1* and *BRCA2*. **(b)** Distribution of type of secondary mutations identified in *BRCA2* depending on whether they were detected on platinum (left pie chart) or PARPi (right pie chart) progression. **(c)** Same as in **b** but in *BRCA1*. **(d)** Distribution of type of secondary mutations on the domain structure of BRCA1 depending on the position of the primary mutation harboured by the patient. **(e)** Length in base pairs of all reversion deletions in BRCA1, assigned to each of the protein domains. **(f)** Same as in **d** but for BRCA2. **(g)** Same as in **e** but for BRCA2. SNV: single-nucleotide variant. DELIN: deletion + insertion. Each square/dot represents a patient. Squares with more than one colour reflect different types of secondary mutations identified in the same patient.

Regarding the type of secondary mutations affecting the different domains of BRCA proteins, deletions of more than 1 bp concentrated in the exon 11 region of BRCA1 (**Fig 2d; Supp Table S2**), as expected by the accumulation of substantial deletion reversions in that area (**Fig 2e**) and the *in vitro* data showing that most of the exon 11 sequence is not required for the function of BRCA1 in generating resistance[43]. A more homogenous distribution was observed in the case of BRCA2 (**Fig 2f**), probably due to the fact that most secondary mutations in this gene were deletions of more than 1 bp (**Fig 2a**). Interestingly, however, the biggest deletions causing reversions in BRCA2 affected the BRC repeats, again suggesting that maintaining only a subset of these could be sufficient to generate resistance (**Fig 2g; Supp Table S2;** see above).

The analysis of secondary mutation types depending on the specific locations of primary mutations showed that mutations affecting the 185delAG region (c.68_69delAG; E23Vfs*) in the RING domain of BRCA1 tend to be small deletions (1 bp; 3 out of 10 cases from 9 patients) or insertions (2 bp; 6 out of 10 cases from 9 patients), similar to what is observed in the 5382insC region (c.5266insC; Q1756Pfs*) in the BRCT domains of BRCA1, with 1 bp deletions (3 out of 5 cases from 4 patients), 2 bp insertions (1 out of 5 cases from 4 patients) and a DELIN (4 bp deletion + 3 bp insertion) accounting for all cases described (**Fig 2d; Supp Table S2**), in line with these domains not being particularly permissive to drastic amino acid changes (**Fig 1f, 2e**). Secondary mutations around the *BRCA2* 6174delT (c.5946delT; S1982Rfs*) region showed more variety, with small insertions (3 out of 13 cases from 5 patients) and deletions ranging from 8 to 137 bp (7 out of 13 cases from 5 patients) (**Fig 2f; Supp Table S2**), as expected due to the flexibility observed in the exon 11 region of the gene (**Fig 1g, 2g**).

In cases of primary mutations caused by SNVs, a clearer correlation between mutation location and mutational mechanisms driving reversion acquisition could be established. For example, the *BRCA1* c.188T>A (L63*) mutation in the RING domain was always reverted through a secondary SNV (2/2 cases), while the *BRCA1* c.1045G>T (E349*) mutation in the exon 11 region was reverted through a secondary SNV (5/7 cases) or a deletion removing the STOP codon (2/7 cases) (**Fig 2d; Supp Table S3**). Particularly interesting is the *BRCA2* primary mutation c.5614A>T (K1872*), which was always reverted though deletions ranging from 27-619 bp (7/7 cases; **Fig 2f; Supp Table S3**). This is in line with the previous observations suggesting less flexibility for amino acid changes in the BRCA1 RING domain, compared to the exon 11 region of both BRCA1 and BRCA2 (see above).

Collectively, these data suggest that both location and nature of the primary mutation can determine to some extent the type of repair event leading to the acquisition of secondary mutations, even in events detected in different patients and on different treatments.

### Secondary mutations involving large deletions have features of microhomology-mediated end-joining (MMEJ) repair

Deletions accounted for the majority of secondary mutations studied, both in *BRCA1* (58.7%) and *BRCA2* (77.6%) (**Fig 2a**). Most deletion events can be explained by the use of error-prone DNA repair mechanisms of end joining, usually classical non-homologous end joining (NHEJ) or alternative end-joining (alt-EJ). A particular sub-pathway of alt-EJ makes use of small sequence microhomologies surrounding the break point, and consequently has been named microhomology-mediated end-joining (MMEJ)[50]. The analysis of microhomologies surrounding secondary deletions detected them in 70.9% cases, and 50.6% involved microhomologies of more than 1 bp, suggestive of MMEJ repair[51] (**Fig 3a; Supp Table S2**).

**Figure 3.**
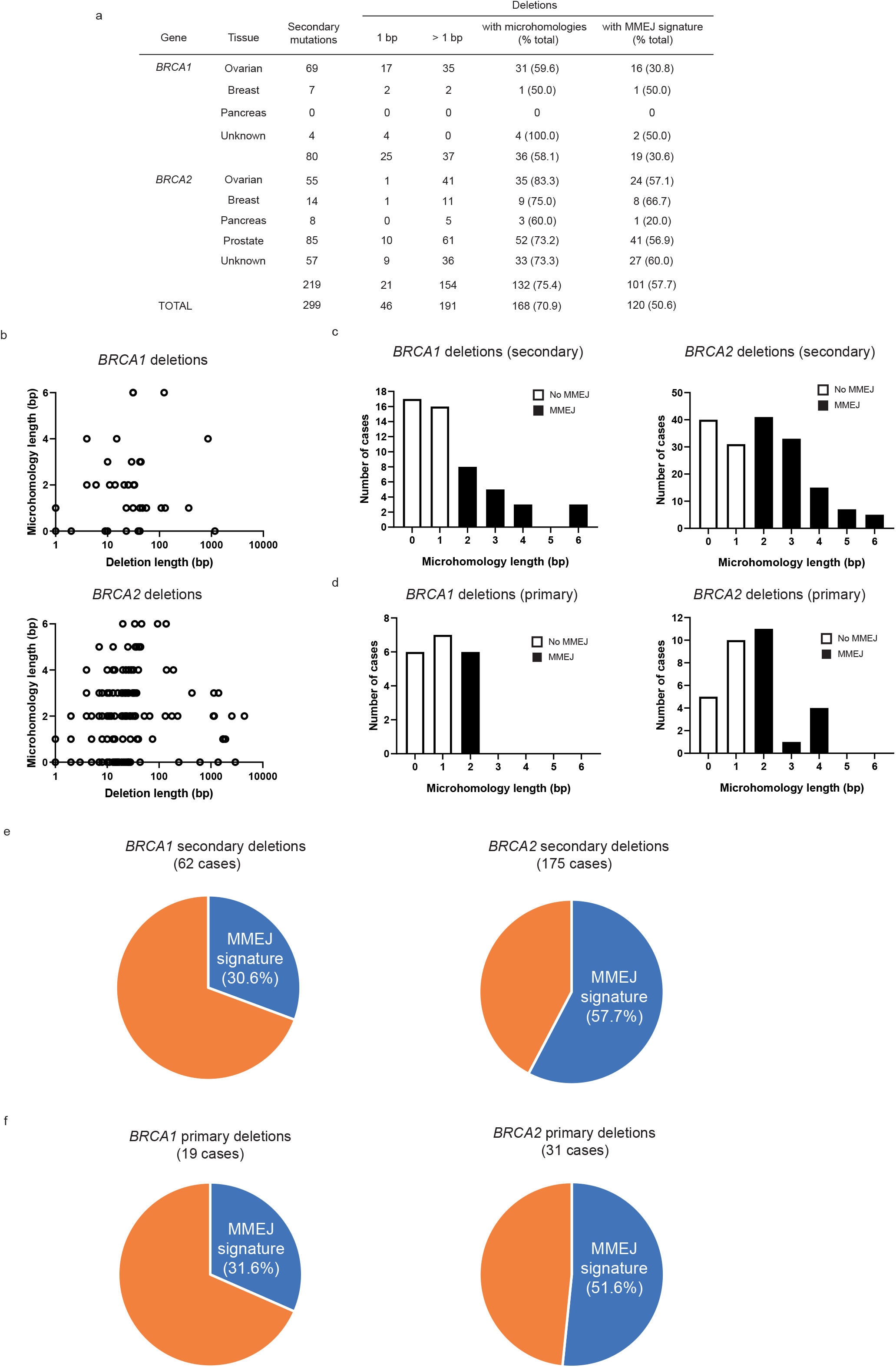
**(a)** Type of deletions and microhomology usage in secondary mutations identified in this patient cohort. **(b)** Lack of correlation between deletion length and microhomology usage in *BRCA1* (top panel) and *BRCA2* (bottom panel) secondary deletions. **(c)** Distribution of microhomology usage in *BRCA1* (left panel) and *BRCA2* (right panel) secondary deletions. **(d)** Distribution of microhomology usage in *BRCA1* (left panel) and *BRCA2* (right panel) primary deletions. **(e)** Prevalence of MMEJ signatures in *BRCA1* (left pie chart) and *BRCA2* (right pie chart) secondary mutations. **(f)** Prevalence of MMEJ signatures in *BRCA1* (left pie chart) and *BRCA2* (right pie chart) primary mutations. MMEJ: microhomology-mediated end joining.

The cases of deletions of more than 1 bp ranged from small deletions of 2 bp to large ones involving as many as almost 3000 bp. The number of base pairs of microhomology detected around breakpoints ranged from 2 to 6 bp (**Supp Table S2**), with no apparent correlation observed between deletion and microhomology lengths (**Fig 3b**). Microhomology lengths in secondary deletions were distributed similarly in *BRCA1* and *BRCA2* cases (**Fig 3c**). Interestingly, we observed an increase in microhomology length usage in secondary mutations in *BRCA* genes (**Fig 3c; Supp Table S2**) when compared to primary mutations (**Fig 3d; Supp Table S3**), although this was not statistically significant (two-tailed two-proportions test, *p* value NS). Importantly, deletions in *BRCA2* were significantly enriched in MMEJ signatures compared to those in *BRCA1*, regardless of their origin being primary or secondary (two-tailed two-proportions test *p* value = 0.0001; **Fig 3e,f; Supp Table S2 and S3**). These results are in agreement with previous reports suggesting that loss of HRR capacity in BRCA deficient tumours upregulates the use of alternative repair pathways, like MMEJ[52].

## Discussion

In this report, we have carried out a retrospective analysis of secondary mutations acquired in tumour and ctDNA samples from patients with mutations in *BRCA1* or *BRCA2* genes and on progression after treatment with platinum or PARPi. Analysis of this cohort, where ovarian cancer was the most represented disease type, detected reversion mutations in 26.3% of cases (**Fig 1a**). This is probably an overestimation of the frequency of reversions in *BRCA* mutant tumours, as several reported in this study come from case report examples or from cohorts with very limited patient numbers (**Supp Table S1**). However, it is also important to note that secondary mutations not directly restoring the open reading frame (and hence not classified as reversions in our analysis; 10 in *BRCA1* mutant patients, 22 in *BRCA2*; **Supp Table S2**), could still allow regaining of protein function through alternative mechanisms. Notwithstanding, *BRCA* reversions are the only confirmed mechanism of resistance identified in clinical samples and their exact prevalence will be better defined with the acquisition of more clinical data from patients progressing on platinum drugs or PARPi.

It is interesting to note that, although it did not reach significance, reversions in *BRCA2* seemed to be more prevalent than in *BRCA1* (**Fig 1a**). This has not translated into patients with *BRCA2* mutated tumours responding worse to treatment, however, as rather the opposite has been observed in some cases[3]. In addition, other mechanisms of resistance may operate more frequently in *BRCA1* mutant tumours, as suggested by pre-clinical studies identifying a wider variety of resistance mechanisms in this genetic background[9]. Whatever the case, it is significant that in both *BRCA1* and *BRCA2* reversions the putative proteins that could be expressed can lack several hundred amino acids (**Fig 1f,g**). The ability of BRCA1 hypomorphic proteins to confer resistance to drug treatment is well documented *in vitro*[43, 53, 54] and has also been recently described in patient-derived xenograft models[19, 55, 56]. Importantly, it has been shown that the protein domains of BRCA1 and BRCA2 play different functions in the roles these proteins have in preserving genomic stability[57–60], suggesting that PARPi-resistant tumours expressing hypomorphic forms of BRCA proteins could be treated with combinations of other targeted agents. For example, the TR2 domain of BRCA2 or isomerization of the RING domain of BRCA1 are required for their function in DNA replication fork protection, but not for HRR[60, 61], which would suggest that reversions affecting the functionality of these regions could be targeted by agents causing replication stress. Some ongoing clinical trials where the PARPi olaparib is being combined with inhibitors of the replication stress response pathway, most notably of the checkpoint kinase ATR (NCT03462342, NCT02576444, NCT04239014, NCT03330847)[62], will provide clinical data where to explore this hypothesis. Our data also suggest that in a post-PARPi treatment scenario, understanding the molecular events leading to reversions could help identifying the best treatment options going forward.

It was surprising to see that the prevalence of type of secondary mutation is different between *BRCA1* and *BRCA2*. Although in both cases deletions were the most frequent event, *BRCA2* secondary mutations are significantly more enriched for deletions, especially of more than 1 bp (**Fig 2a**). Although it did not reach statistical significance, a similar trend was also observed when analysing the primary mutations carried from the germline in this patient cohort (**Fig 1c; Supp Table S3,** two-tailed two-proportions test *p* value=NS), which could suggest that variables such as chromosomal location and/or chromatin landscape around *BRCA1* and *BRCA2* genes could be important in determining the repair pathways at play when genetic alterations occur in these genes[63]. Although we did not focus on mutational mechanisms driving reversions through SNVs due to the smaller number of cases, it will be interesting to understand the contribution of other DNA repair pathways such as translesion synthesis, nucleotide excision repair, base excision repair or mismatch repair to such outcomes.

The prevalence of deletions as the main mechanism of acquisition of reversion mutations in *BRCA* genes suggests the presence of a DNA double-strand break intermediate that is repaired by end-joining mechanisms. Mutational signatures of NHEJ usually involve generation of small INDELs by limited DNA-end resection imposed by the presence of the Ku heterodimer bound to the ends of the DNA break. On the other hand, signatures of MMEJ repair involve more extensive DNA-end resection and the use of microhomologies (2-6 bp) flanking the break site. These can be placed several hundred base pairs apart, which can lead to sizable deletions and chromosomal translocations[50]. Strikingly, we observed significant microhomology usage in deletions affecting *BRCA* genes, suggestive of prevalence of MMEJ repair mechanisms, especially in *BRCA2* (**Fig 3c-e**). A key player in MMEJ repair is DNA polymerase theta, encoded by the *POLQ* gene[64], which has been shown to be essential for cell survival in *BRCA*-deficient cell lines and to compete with HRR proteins for similar DNA repair substrates[52, 65]. Compounds inhibiting DNA-PK, the key protein kinase involved in NHEJ, are entering the clinic[66, 67]. It will be interesting to test whether blockage of NHEJ and/or MMEJ repair in *BRCA* mutant backgrounds has the potential to prevent the accumulation of reversion mutations, and hence the appearance of resistance to drug treatments.

## Acknowledgements

The authors would like to thank Dr Jessica Brown and Dr Natasha Lukashchuk (AstraZeneca) for critical reading of the manuscript and provision of insightful comments.

## Methods

Primary and secondary mutation data from *BRCA* genes were collected from the literature and formatted in the standard coding DNA reference sequence (**Supp Table S2** and references therein). For the analysis of types of secondary mutations, we analysed the presence of microhomologies in deletions by following the strategy described by Taheri-Ghahfarokhi et al[51]. In short, when the mutation was a pure deletion, we first located the deleted sequence in the full gene sequence and adjusted its position if the nucleotides before the deletion were the same as the last nucleotides of the deleted sequence. Then, we checked how many contiguous nucleotides at the beginning of the deleted sequence could be matched after the deletion. The number of contiguously matched nucleotides is equal to the length of the microhomology, with a maximum length equal to the length of the deletion. Microhomologies of at least 2 bp were considered compatible with MMEJ repair (see example below).

**Figure caption.**
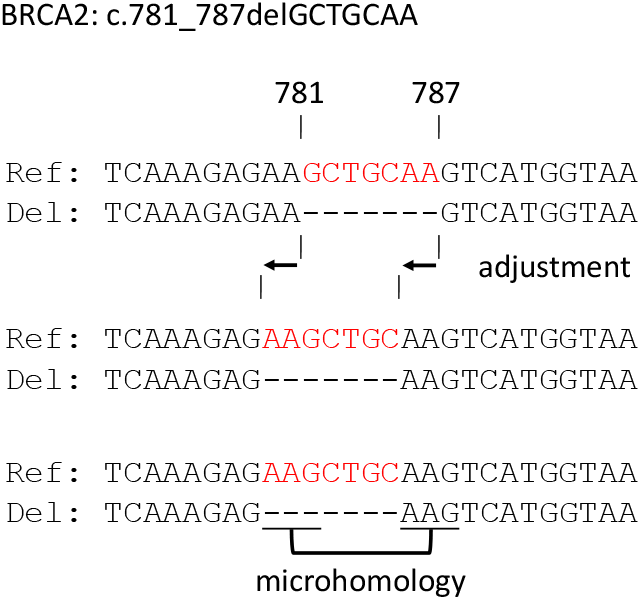
Procedure for microhomology identification. First, the reported deletion is matched to the reference sequence. If necessary, the position of the deletion is adjusted. In this example, reporting the deletion as c.781_787del or c.779_785del would result in the same deleted sequence, so we redefined the position in which the deletion happened. Finally, the length of the microhomology is determined by comparing the nucleotides at the beginning of the deleted sequence with those after the deletion. Without the adjustment procedure, only a microhomology of length 2 would have been reported, when a microhomology of length 3 could be reported.

The analysis was done in R (https://www.R-project.org/) version 3.6.0 and sequences were retrieved using the biomaRt package[68, 69]. We used the reference sequences NM_007294 for *BRCA1* and NM_000059 for *BRCA2*. For cases in which the break points of the secondary mutation fell very close to the primary mutation, we checked the outcome manually with the help of Mutalyzer[70].

A test of Equal or Given Proportions was used to assess if an observed difference in proportion between two groups is statistically significant. The analysis was done in R (https://www.R-project.org/) version 3.6.0 using the prop.test function.

## Supplementary Material

**Supplementary Table S1.** Data sources

**Supplementary Table S2.** Primary and secondary mutations in *BRCA* genes

**Supplementary Table S3.** Type and origin of primary mutations and reversions in *BRCA* genes

**Supplementary Table S4.** Reversion mutations in *BRCA2* identified *in vitro*

## References

1. Nielsen FC, van Overeem Hansen T, Sorensen CS. Hereditary breast and ovarian cancer: new genes in confined pathways. Nat Rev Cancer 2016; 16: 599–612.

2. Chen CC, Feng W, Lim PX et al. Homology-Directed Repair and the Role of BRCA1, BRCA2, and Related Proteins in Genome Integrity and Cancer. Annu Rev Cancer Biol 2018; 2: 313–336.

3. Hollis RL, Churchman M, Gourley C. Distinct implications of different BRCA mutations: efficacy of cytotoxic chemotherapy, PARP inhibition and clinical outcome in ovarian cancer. Onco Targets Ther 2017; 10: 2539–2551.

4. Farmer H, McCabe N, Lord CJ et al. Targeting the DNA repair defect in BRCA mutant cells as a therapeutic strategy. Nature 2005; 434: 917.

5. Bryant HE, Schultz N, Thomas HD et al. Specific killing of BRCA2-deficient tumours with inhibitors of poly(ADP-ribose) polymerase. Nature 2005; 434: 913.

6. Yap TA, Plummer R, Azad NS, Helleday T. The DNA Damaging Revolution: PARP Inhibitors and Beyond. Dev Therap Tumor Biol 2019; 185–195.

7. Golan T, Hammel P, Reni M et al. Maintenance Olaparib for Germline BRCA-Mutated Metastatic Pancreatic Cancer. N Engl J Med 2019; 381: 317–327.

8. de Bono J, Mateo J, Fizazi K et al. Olaparib for Metastatic Castration-Resistant Prostate Cancer. N Engl J Med 2020; 382: 2091–2102.

9. D’Andrea AD. Mechanisms of PARP inhibitor sensitivity and resistance. DNA Repair 2018; 71: 172–176.

10. Edwards SL, Brough R, Lord CJ et al. Resistance to therapy caused by intragenic deletion in BRCA2. Nature 2008; 451: 1111.

11. Sakai W, Swisher EM, Karlan BY et al. Secondary mutations as a mechanism of cisplatin resistance in BRCA2-mutated cancers. Nature 2008; 451: 1116.

12. Swisher EM, Sakai W, Karlan BY et al. Secondary BRCA1 Mutations in BRCA1-Mutated Ovarian Carcinomas with Platinum Resistance. Cancer Res 2008; 68: 2581–2586.

13. Weigelt B, Comino-Méndez I, de Bruijn I et al. Diverse BRCA1 and BRCA2 Reversion Mutations in Circulating Cell-Free DNA of Therapy-Resistant Breast or Ovarian Cancer. Clin Cancer Res 2017; 23: 6708–6720.

14. Afghahi A, Timms KM, Vinayak S et al. Tumor BRCA1 Reversion Mutation Arising during Neoadjuvant Platinum-Based Chemotherapy in Triple-Negative Breast Cancer Is Associated with Therapy Resistance. Clin Cancer Res 2017; 23: 3365–3370.

15. Gornstein EL, Sandefur S, Chung JH et al. BRCA2 Reversion Mutation Associated With Acquired Resistance to Olaparib in Estrogen Receptor-positive Breast Cancer Detected by Genomic Profiling of Tissue and Liquid Biopsy. Clinical Breast Cancer 2018; 18: 184–188.

16. Waks AG, Cohen O, Kochupurakkal B et al. Reversion and non-reversion mechanisms of resistance to PARP inhibitor or platinum chemotherapy in BRCA1/2-mutant metastatic breast cancer. Annals of Oncology 2020.

17. Yap TA, Kristeleit R, Michalarea V et al. Phase I trial of the poly(ADP-ribose) polymerase (PARP) inhibitor olaparib and AKT inhibitor capivasertib in patients with BRCA1/2 and non-BRCA1/2 mutant cancers. Cancer Discov 2020; CD-20–0163.

18. Meijer TG, Verkaik NS, Deurzen CHMv et al. Direct Ex Vivo Observation of Homologous Recombination Defect Reversal After DNA-Damaging Chemotherapy in Patients With Metastatic Breast Cancer. JCO Precis Oncol 2019; 1–12.

19. Cruz C, Castroviejo-Bermejo M, Gutiérrez-Enríquez S et al. RAD51 foci as a functional biomarker of homologous recombination repair and PARP inhibitor resistance in germline BRCA-mutated breast cancer. Annals of Oncology 2018; mdy099–mdy099.

20. Barber LJ, Sandhu S, Chen L et al. Secondary mutations in BRCA2 associated with clinical resistance to a PARP inhibitor. J Pathol 2013; 229: 422–429.

21. Banda K, Swisher EM, Wu D et al. Somatic Reversion of Germline BRCA2 Mutation Confers Resistance to Poly(ADP-ribose) Polymerase Inhibitor Therapy. J Clin Oncol Precis Oncol 2018; 1–6.

22. Norquist B, Wurz KA, Pennil CC et al. Secondary Somatic Mutations Restoring BRCA1/2 Predict Chemotherapy Resistance in Hereditary Ovarian Carcinomas. J Clin Oncol 2011; 29: 3008–3015.

23. Christie EL, Fereday S, Doig K et al. Reversion of BRCA1/2 Germline Mutations Detected in Circulating Tumor DNA From Patients With High-Grade Serous Ovarian Cancer. J Clin Oncol 2017; 35: 1274–1280.

24. Kondrashova O, Nguyen M, Shield-Artin K et al. Secondary Somatic Mutations Restoring RAD51C and RAD51D Associated with Acquired Resistance to the PARP Inhibitor Rucaparib in High-Grade Ovarian Carcinoma. Cancer Discovery 2017; 7: 984–998.

25. Lin KK, Harrell MI, Oza AM et al. BRCA Reversion Mutations in Circulating Tumor DNA Predict Primary and Acquired Resistance to the PARP Inhibitor Rucaparib in High-Grade Ovarian Carcinoma. Cancer Discov 2019; 9: 210–219.

26. Patch A-M, Christie EL, Etemadmoghadam D et al. Whole–genome characterization of chemoresistant ovarian cancer. Nature 2015; 521: 489.

27. Mayor P, Gay LM, Lele S, Elvin JA. BRCA1 reversion mutation acquired after treatment identified by liquid biopsy. Gynecologic Oncology Reports 2017; 21: 57–60.

28. Patel JN, Braicu I, Timms KM et al. Characterisation of homologous recombination deficiency in paired primary and recurrent high-grade serous ovarian cancer. British Journal of Cancer 2018; 119: 1060–1066.

29. Jacob SL, Kiedrowski LA, Chae YK. The dynamic landscape of BRCA1 reversion mutations from indel to SNV in a patient with ovarian cancer treated with PARP-inhibitors and immunotherapy. Heliyon 2020; 6: e03841.

30. Khalique S, Pettitt SJ, Kelly G et al. Longitudinal analysis of a secondary BRCA2 mutation using digital droplet PCR. J Pathol 2020; 6: 3–11.

31. Shroff RT, Hendifar A, McWilliams RR et al. Rucaparib Monotherapy in Patients With Pancreatic Cancer and a Known Deleterious BRCA Mutation. JCO Precis Oncol 2018; 1–15.

32. Tao H, Liu S, Huang D et al. Acquired multiple secondary BRCA2 mutations upon PARPi resistance in a metastatic pancreatic cancer patient harboring a BRCA2 germline mutation. American journal of translational research 2020; 12: 612–617.

33. Pishvaian MJ, Biankin AV, Bailey P et al. BRCA2 secondary mutation-mediated resistance to platinum and PARP inhibitor-based therapy in pancreatic cancer. British Journal of Cancer 2017; 116: 1021–1026.

34. Quigley D, Alumkal JJ, Wyatt AW et al. Analysis of Circulating Cell-Free DNA Identifies Multiclonal Heterogeneity of BRCA2 Reversion Mutations Associated with Resistance to PARP Inhibitors. Cancer Discovery 2017; 7: 999–1005.

35. Goodall J, Mateo J, Yuan W et al. Circulating Cell-Free DNA to Guide Prostate Cancer Treatment with PARP Inhibition. Cancer Discovery 2017; 7: 1006–1017.

36. Simmons AD, Nguyen M, Pintus E. Polyclonal BRCA2 mutations following carboplatin treatment confer resistance to the PARP inhibitor rucaparib in a patient with mCRPC: a case report. BMC Cancer 2020; 20: 215.

37. Carneiro BA, Collier KA, Nagy RJ et al. Acquired Resistance to Poly (ADP-ribose) Polymerase Inhibitor Olaparib in BRCA2-Associated Prostate Cancer Resulting From Biallelic BRCA2 Reversion Mutations Restores Both Germline and Somatic Loss-of-Function Mutations. JCO Precis Oncol 2018; 1–8.

38. Cheng HH, Salipante SJ, Nelson PS et al. Polyclonal BRCA2 Reversion Mutations Detected in Circulating Tumor DNA After Platinum Chemotherapy in a Patient With Metastatic Prostate Cancer. JCO Precis Oncol 2018; 1–5.

39. Vidula N, Rich TA, Sartor O et al. Routine plasma-based genotyping to comprehensively detect germline, somatic, and reversion BRCA mutations among patients with advanced solid tumors. Clin Cancer Res 2020; clincanres.2933.2019.

40. Rebbeck TR, Mitra N, Wan F et al. Association of type and location of BRCA1 and BRCA2 mutations with risk of breast and ovarian cancer. JAMA 2015; 313: 1347–1361.

41. Moore K, Colombo N, Scambia G et al. Maintenance Olaparib in Patients with Newly Diagnosed Advanced Ovarian Cancer. N Engl J Med 2018; 379: 2495–2505.

42. Levy-Lahad E, Catane R, Eisenberg S et al. Founder BRCA1 and BRCA2 mutations in Ashkenazi Jews in Israel: frequency and differential penetrance in ovarian cancer and in breast-ovarian cancer families. Am J Human Genet 1997; 60: 1059–1067.

43. Wang Y, Bernhardy AJ, Cruz C et al. The BRCA1-Delta11q Alternative Splice Isoform Bypasses Germline Mutations and Promotes Therapeutic Resistance to PARP Inhibition and Cisplatin. Cancer Res 2016; 76: 2778–2790.

44. Densham RM, Garvin AJ, Stone HR et al. Human BRCA1-BARD1 ubiquitin ligase activity counteracts chromatin barriers to DNA resection. Nat Struct Mol Biol 2016; 23: 647–655.

45. Findlay GM, Daza RM, Martin B et al. Accurate classification of BRCA1 variants with saturation genome editing. Nature 2018; 562: 217–222.

46. Anantha RW, Simhadri S, Foo TK et al. Functional and mutational landscapes of BRCA1 for homology-directed repair and therapy resistance. Elife 2017; 6.

47. Pellegrini L, Yu DS, Lo T et al. Insights into DNA recombination from the structure of a RAD51-BRCA2 complex. Nature 2002; 420: 287–293.

48. Siaud N, Barbera MA, Egashira A et al. Plasticity of BRCA2 Function in Homologous Recombination: Genetic Interactions of the PALB2 and DNA Binding Domains. PLOS Genetics 2011; 7: e1002409.

49. Jonkers J, Meuwissen R, van der Gulden H et al. Synergistic tumor suppressor activity of BRCA2 and p53 in a conditional mouse model for breast cancer. Nature Genetics 2001; 29: 418–425.

50. Chang HHY, Pannunzio NR, Adachi N, Lieber MR. Non-homologous DNA end joining and alternative pathways to double-strand break repair. Nature Reviews Molecular Cell Biology 2017; 18: 495.

51. Taheri-Ghahfarokhi A, Taylor BJ M, Nitsch R et al. Decoding non-random mutational signatures at Cas9 targeted sites. Nucleic Acids Research 2018; 46: 8417–8434.

52. Ceccaldi R, Liu JC, Amunugama R et al. Homologous-recombination-deficient tumours are dependent on Poltheta-mediated repair. Nature 2015; 518: 258–262.

53. Johnson N, Johnson SF, Yao W et al. Stabilization of mutant BRCA1 protein confers PARP inhibitor and platinum resistance. Proceedings of the National Academy of Sciences 2013; 110: 17041–17046.

54. Drost R, Dhillon KK, van der Gulden H et al. BRCA1185delAG tumors may acquire therapy resistance through expression of RING-less BRCA1. J Clin Invest 2016; 126: 2903–2918.

55. Castroviejo-Bermejo M, Cruz C, Llop-Guevara A et al. A RAD51 assay feasible in routine tumor samples calls PARP inhibitor response beyond BRCA mutation. EMBO Molecular Medicine 2018.

56. Wang Y, Bernhardy AJ, Nacson J et al. BRCA1 intronic Alu elements drive gene rearrangements and PARP inhibitor resistance. Nature Communications 2019; 10: 5661.

57. Shakya R, Reid LJ, Reczek CR et al. BRCA1 tumor suppression depends on BRCT phosphoprotein binding, but not its E3 ligase activity. Science 2011; 334: 525–528.

58. Drost R, Bouwman P, Rottenberg S et al. BRCA1 RING function is essential for tumor suppression but dispensable for therapy resistance. Cancer Cell 2011; 20: 797–809.

59. Zhu Q, Pao GM, Huynh AM et al. BRCA1 tumour suppression occurs via heterochromatin-mediated silencing. Nature 2011; 477: 179–184.

60. Schlacher K, Christ N, Siaud N et al. Double-strand break repair-independent role for BRCA2 in blocking stalled replication fork degradation by MRE11. Cell 2011; 145: 529–542.

61. Daza-Martin M, Starowicz K, Jamshad M et al. Isomerization of BRCA1–BARD1 promotes replication fork protection. Nature 2019; 571: 521–527.

62. Forment JV, O’Connor MJ. Targeting the replication stress response in cancer. Pharmacol Ther 2018; 188: 155–167.

63. Clouaire T, Rocher V, Lashgari A et al. Comprehensive Mapping of Histone Modifications at DNA Double-Strand Breaks Deciphers Repair Pathway Chromatin Signatures. Molecular Cell 2018; 72: 250–262.e256.

64. Wyatt DW, Feng W, Conlin MP et al. Essential Roles for Polymerase theta-Mediated End Joining in the Repair of Chromosome Breaks. Mol Cell 2016; 63: 662–673.

65. Mateos-Gomez PA, Gong F, Nair N et al. Mammalian polymerase theta promotes alternative NHEJ and suppresses recombination. Nature 2015; 518: 254–257.

66. Harnor SJ, Brennan A, Cano C. Targeting DNA-Dependent Protein Kinase for Cancer Therapy. ChemMedChem 2017; 12: 895–900.

67. Fok JHL, Ramos-Montoya A, Vazquez-Chantada M et al. AZD7648 is a potent and selective DNA-PK inhibitor that enhances radiation, chemotherapy and olaparib activity. Nature Communications 2019; 10: 5065.

68. Durinck S, Moreau Y, Kasprzyk A et al. BioMart and Bioconductor: a powerful link between biological databases and microarray data analysis. Bioinformatics 2005; 21: 3439–3440.

69. Durinck S, Spellman PT, Birney E, Huber W. Mapping identifiers for the integration of genomic datasets with the R/Bioconductor package biomaRt. Nature Protocols 2009; 4: 1184–1191.

70. Wildeman M, van Ophuizen E, den Dunnen JT, Taschner PEM. Improving sequence variant descriptions in mutation databases and literature using the Mutalyzer sequence variation nomenclature checker. Hum Mutat 2008; 29: 6–13.

